# Loss of macrodomain catalytic activity modulates Chikungunya virus dissemination and transmission potential in *Aedes* mosquitoes

**DOI:** 10.1101/2025.10.09.681498

**Authors:** Eugenia S. Bardossy, Lena Bergmann, Annabelle Henrion-Lacritick, Jared Nigg, Galen J. Correy, Alan Ashworth, James S. Fraser, Maria-Carla Saleh

## Abstract

Viral macrodomains are promising antiviral targets that counteract host ADP-ribosylation-mediated antiviral responses in mammals. However, their role in dual-host viruses within the mosquito vector remains largely unknown. Here, we investigated the role of the Chikungunya virus (CHIKV) nsP3 macrodomain catalytic activity by mutating the active-site asparagine 24 (N24). N24 mutant viruses rapidly selected for compensatory second-site mutations at aspartate 31 (D31) in mammalian Vero cells, while interferon-competent human cells favored reversion at position 24 instead, suggesting adaptive responses to macrodomain functional impairment. Catalytically inactive double-mutant viruses replicated poorly in interferon-competent human cells but comparably to wild-type in mosquito cells. In *Aedes* mosquitoes *in vivo*, macrodomain mutations modulated CHIKV replication and infectivity in a mutation-specific manner: one mutant reduced them while another enhanced them. Biochemical and structural analyses confirmed that these phenotypes arise in the context of a catalytically inactive macrodomain that retains ADP-ribose binding capacity, with D31 mutations lining the substrate exit path. These findings demonstrate that macrodomain catalytic activity modulates CHIKV dissemination and transmission potential in *Aedes* mosquitoes, revealing fundamental differences in host-dependent evolutionary advantage for a catalytically inactive viral enzyme.

## Introduction

The emergence and re-emergence of RNA viruses pose ongoing global public health threats, from pandemic coronaviruses to mosquito-borne arboviruses such as chikungunya, dengue, and Zika. Identifying conserved viral proteins tractable to therapeutic targeting is therefore a priority. Viral macrodomains are conserved enzymatic domains encoded within the non structural genes of alphaviruses, coronaviruses, rubella virus, and hepatitis E virus^1,2^, that have emerged as attractive antiviral targets^3–6^. These domains counteract host ADP-ribosylation-based antiviral defenses central to the mammalian innate immune response^7–13^, and recent identification of promising small-molecule inhibitors targeting the SARS-CoV-2 macrodomain^14^ has provided proof-of-concept for this therapeutic strategy.

Viral macrodomains exhibit ADP-ribosylhydrolase catalytic activity, binding to mono- and poly-ADP-ribosylated (MARylated and PARylated) proteins and cleaving the ADP-ribose modification, an “eraser” function that directly counters the activity of host poly(ADP-ribose) polymerases (PARPs)^7–12^. Stimulated by interferon (IFN) signaling, PARPs catalyze the addition of ADP-ribose moieties to viral and host target proteins as part of the antiviral response^15–21^, and viral macrodomains neutralize this defense by reversing these modifications^22–24^. Beyond catalytic erasure, macrodomains can also function as “readers”, binding ADP-ribosylation marks to facilitate protein interactions or to direct modified substrates toward other functional domains^25^. For viral macrodomains, the relative contribution of eraser versus reader activity to viral fitness has not been fully resolved and may critically depend on host context^9^.

This question is particularly relevant for dual-host arboviruses such as chikungunya virus (CHIKV), which must replicate alternately in mammalian vertebrate hosts and *Aedes* mosquito vectors, environments with fundamentally different immune systems and physiological conditions. In mammals, IFN signaling and PARP-mediated ADP-ribosylation are central components of the antiviral response, making macrodomain eraser activity functionally critical^4,7,10–12,23,24,26^. In mosquitoes, antiviral immunity is organized primarily around RNA interference rather than IFN signaling^27–30^, raising the fundamental question of whether macrodomain catalytic activity serves the same function, or any essential function, in the insect host. Despite the importance of this question for both basic virology and antiviral drug development, the role of viral macrodomains in the mosquito vector has remained almost entirely unexplored.

Comparative genomics provides an important clue. The catalytic asparagine at position 24 (N24), essential for ADP-ribosylhydrolase activity^4,12,31–34^, is invariably conserved across dual-host alphaviruses and other human-pathogenic viruses, but is consistently absent in mosquito-specific alphaviruses that are incapable of infecting vertebrates^35,36^. This recurrent loss strongly implies that the selective pressure maintaining macrodomain catalytic activity originates from the vertebrate host, while this pressure is relaxed in viruses replicating exclusively in mosquitoes. These observations raise the question of whether a dual-host alphavirus such as CHIKV could tolerate the loss of macrodomain catalytic activity in mosquitoes, and whether the macrodomain’s reader function might be sufficient to support viral replication in the absence of catalysis.

CHIKV is an ideal model to address these questions. A re-emergent human pathogen responsible for millions of cases of febrile arthralgia worldwide^37^, CHIKV is transmitted by *Aedes albopictus* and *Aedes aegypti* mosquitoes and has undergone substantial global expansion in recent decades, with ongoing outbreaks in tropical and subtropical regions. Its positive-sense RNA genome encodes four non-structural proteins (nsP1–nsP4) forming the viral replicase complex, and the structural proteins required for virion assembly^37^. Among these, non-structural protein 3 (nsP3) is a determinant of mosquito vector specificity in CHIKV^38^, a role also described for the related O’nyong-nyong virus^39^, and of neurovirulence in Semliki Forest and Sindbis viruses^40–43^. In CHIKV, nsP3 also harbors the catalytically active macrodomain within its N-terminal 160 amino acids, whose ADP-ribosylhydrolase activity is well characterized^9,11,12,33,34^. However, its function in the mosquito has not been examined *in vivo*.

Here, we investigated the role of the CHIKV nsP3 macrodomain during viral replication in mammalian and mosquito hosts. Using CHIKV as a model, we engineered recombinant viruses carrying mutations in the macrodomain that eliminate its catalytic activity. Attempts to propagate these viruses rapidly selected for second-site mutations at residue D31, which did not restore catalytic activity. The infection of mammalian cells and laboratory colonies of *Aedes* mosquitoes with wild-type and mutant viruses reveals unexpected and host-dependent effects of macrodomain catalytic inactivation on viral fitness and demonstrates that the selective pressures acting on this domain differ fundamentally between mammalian and mosquito hosts.

## Results

### N24 mutations in the CHIKV macrodomain catalytic site rapidly select for second-site mutations at residue D31

The asparagine at position 24 (N24) of the CHIKV nsP3 macrodomain is located at the active site and makes direct contacts with bound ADP-ribose^31,34^ (Fig. 1a). Substitution of the equivalent asparagine in the SARS-CoV-2 macrodomain abolishes ADP-ribosylhydrolase catalytic activity^4,11,33,34^, establishing N24 as essential for catalytic function. To investigate the role of macrodomain catalytic activity in CHIKV replication, we introduced two mutations at this position into the infectious clone of the Caribbean CHIKV strain: N24A (asparagine to alanine) and N24D (asparagine to aspartic acid), both of which abolish macrodomain catalytic activity in the homologous SARS-CoV-2 system^4^ (Fig. 1b). We included both mutations because the N24A substitution reduces thermodynamic stability in SARS-CoV-2, whereas N24D abolishes catalytic activity without destabilizing the protein^4^. To produce viral stocks, *in vitro* transcribed RNAs encoding WT, N24A, or N24D viruses were transfected into Vero cells. Cell culture supernatants were collected at one day post-transfection (P0) and used to infect fresh Vero cells. Supernatants from this passage were collected at one and two days post-infection (P1). Total RNA was extracted from supernatants at each timepoint, the N-terminal region of the nsP3 macrodomain (covering the first 119 amino acids) was amplified by RT-PCR, and the products were analyzed by Sanger sequencing (Fig. 1b). This region was selected to confirm the stability of the introduced mutations at position 24 and to monitor the emergence of potential second-site mutations in its immediate vicinity.

**Figure 1.**
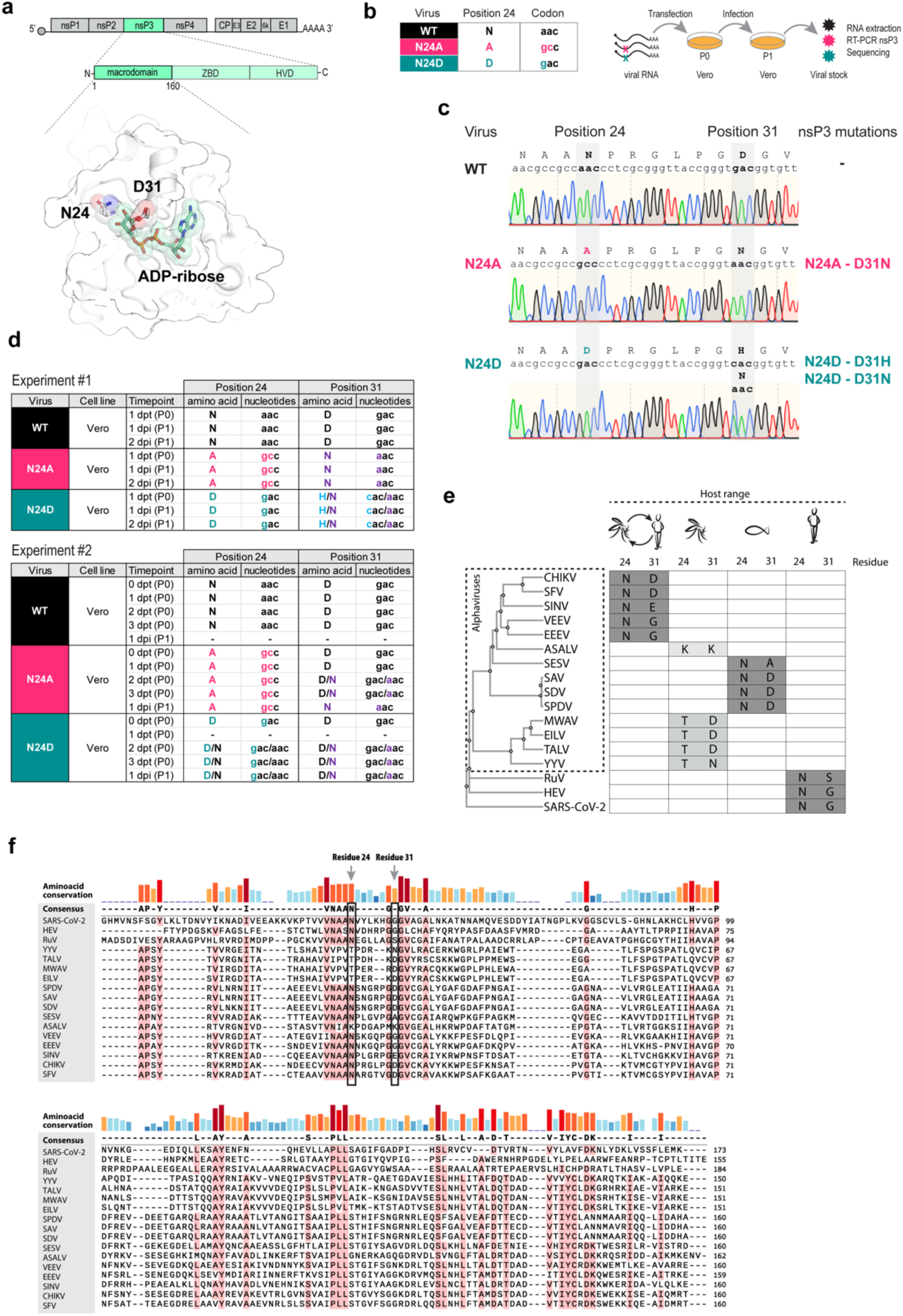
N24 mutations in the CHIKV macrodomain catalytic site rapidly select for second-site mutations at residue D31. a, Schematic of the CHIKV genome highlighting the nsP3 gene. nsP3 comprises an N-terminal macrodomain (first 160 amino acids), a central zinc-binding domain (ZBD), and a C-terminal hypervariable domain (HVD). Below, structure of the CHIKV macrodomain (PDB 3GPO) highlighting the catalytic asparagine 24 (N24), residue 31 and bound ADP-ribose. **b,** N24A (asparagine to alanine) and N24D (asparagine to aspartic acid) mutations were introduced into the CHIKV Caribbean infectious clone. *In vitro* transcribed RNAs were transfected into Vero cells, supernatants were collected at one day post-transfection (P0) and used to infect fresh Vero cells, and supernatants from this passage (P1) were collected at one and two days post-infection. All samples were processed by RNA extraction, RT-PCR of the first 119 amino acids of nsP3 and Sanger sequencing. **c,** Representative Sanger chromatograms from P1 viral stocks. The WT sequence is unchanged at position 24, whereas N24A and N24D viruses carry second-site mutations at residue 31 (D31N, and D31H/N, respectively). **d,** Amino acid and nucleotide identities at residues 24 and 31 in two independent experiments in Vero cells. In Experiment #1, supernatants were collected at one day post-transfection (P0) and one and two days post-infection (P1). In Experiment #2, additional timepoints were sampled at zero to three days post-transfection (P0) and one day post-infection (P1). Dashes indicate timepoints with insufficient viral RNA for sequencing, and ‘/’ denotes mixed populations. **e,** Residues 24 and 31 in macrodomains of 14 alphaviruses grouped by host range: dual-host (CHIKV, SFV, SINV, VEEV, EEEV), mosquito-specific (ASALV, MWAV, EILV, TALV, YYV) and aquatic (SESV, SAV, SDV, SPDV), together with three other human-pathogenic viruses (RuV, HEV, SARS-CoV-2). The phylogram was built using Clustal Omega. Sequence identifiers and databases are listed in Supplementary Table 1. **f,** Multiple sequence alignment of the same 17 macrodomains. Amino acid conservation and the consensus sequence are shown above. Residues conserved in at least 80% of the sequences are highlighted in pink. Arrows mark residues 24 and 31. Amino acid abbreviations: N, asparagine; A, alanine; D, aspartic acid; H, histidine.

Sequencing of the WT virus revealed no changes in the analyzed region of the nsP3 macrodomain at any examined timepoint, confirming genetic stability of the WT sequence under these conditions (Fig. 1c, 1d). In contrast, both N24 mutant viruses rapidly acquired second-site mutations at residue 31 during viral replication, while the introduced mutations at position 24 remained stable. In the N24A mutant, aspartic acid at position 31 was replaced by asparagine (D31N). In the N24D mutant, position 31 acquired either a D31N (GAC to AAC) or D31H (GAC to CAC) substitution, with both variants detected as a mixed population (Fig. 1c, 1d). Consequently, while the N24A mutant yielded a genetically defined double mutant virus (N24A-D31N), the N24D mutant yielded a mixed population of N24D-D31N and N24D-D31H viruses. We confirmed the emergence of D31 mutations in a second independent experiment in Vero cells, in which additional timepoints were sampled (Fig. 1d, Experiment #2). In this second experiment, only the D31N variant was recovered, while D31H was not detected. Notably, for the N24D mutant, partial reversion of N24D towards the WT sequence was observed, and position 31 remained as a mixed population of D31 and D31N variants. In addition, to further confirm the reproducibility of D31 mutation emergence across different cell types, we performed additional viral RNA transfections in BHK-21 mammalian cells and C6/36 mosquito cells (Supplementary Fig. 1a). For the N24A mutant, the D31N mutation emerged in both BHK-21 and C6/36 cells. For the N24D mutant, the D31N mutation emerged only in BHK-21 cells, and partial reversion of N24D to the WT sequence was also observed in this cell line. Taken together, our sequence analyses demonstrate that second-site mutations at residue D31 reproducibly emerge when viral genomes carrying primary mutations at the catalytic N24 site replicate in both mammalian and mosquito cell lines.

Next, we investigated whether the emergence of D31 mutations is a conserved feature across different CHIKV lineages. There are three main CHIKV lineages: African, East-Central-South African (ECSA), and Asian. Our previous experiments were performed with the Caribbean strain from the Asian lineage, which was responsible for the introduction of CHIKV into the Americas. We also introduced N24A and N24D mutations into an infectious clone of the Indian Ocean strain, a representative of the ECSA lineage responsible for the 2005 outbreak in La Réunion. As before, we assessed the genetic stability of N24 mutations and the potential emergence of secondary mutations during viral stock production. Viral RNAs were transfected into BHK-21 cells (P0), passaged once in BHK-21 cells (P1), and then passaged a second time in Vero cells (P2). The D31N secondary mutation consistently emerged in both N24A and N24D Indian Ocean mutant viruses in mammalian cells (Supplementary Fig. 1b). Taken together, our results show that CHIKVs carrying single N24A or N24D mutations are not stably maintained in the mammalian cell lines evaluated, independently of the viral lineage. In this scenario, viral replication is restored by the emergence of compensatory mutations at residue 31 and, in some cases, by partial reversion of the N24 mutation. Notably, in C6/36 mosquito cells the N24D single mutant remained unchanged at both positions throughout the experiment, while the N24A mutant acquired D31N only at passage 1.

To place these mutations in an evolutionary context, we aligned macrodomain sequences from 14 alphaviruses spanning dual-host, aquatic and mosquito-specific host ranges, together with rubella virus, hepatitis E virus and SARS-CoV-2 (Fig. 1e, 1f). Conservation analysis of viral macrodomains shows that, unlike N24, residue 31 is poorly conserved: aspartic acid (D), glutamic acid (E), glycine (G), alanine (A), serine (S), asparagine (N) and lysine (K) are all found at this position across alphaviruses and other viral macrodomains (Fig. 1e, 1f). Notably, histidine (H) is not found at residue 31 in any of the viruses analyzed, and asparagine (N) occurs only in the mosquito-specific Yada Yada virus.

Because a second-site mutation at the equivalent position has been previously reported in Sindbis virus during replication of a macrodomain double mutant^43^, our results suggest that residue 31 represents a recurrent site of genetic compensation across alphaviruses. This raises the question of whether the selective pressure driving D31 second-site mutation emergence operates similarly in immunologically relevant human cells with a competent innate immune response.

### N24 catalytic mutations impair CHIKV replication in interferon-competent human cells but not in mosquito cells

The rapid emergence of D31 second-site mutations during viral stock production in Vero cells suggested that N24 catalytic mutations impose a fitness cost during viral replication. To investigate whether this selective pressure operates in interferon-competent human cells, we transfected N24A and N24D viral RNAs into interferon-competent A549 cells and monitored the genetic stability of the introduced mutations over time (Fig. 2a, 2b). For the N24A mutant, pure N24A sequence was detected at days 2 and 3 post-transfection, with no D31 second-site mutations emerging at any timepoint. At passage 1, a mixture of N24A and WT sequence was detected at position 24, indicating partial reversion toward the WT asparagine. For the N24D mutant, pure N24D sequence was detected at all four P0 timepoints, again with no D31 second-site mutations. At passage 1, a mixture of N24D and WT sequence was detected at position 24, consistent with partial reversion. In neither case did D31 second-site mutations emerge, in contrast to the rapid D31 emergence observed in Vero cells. The low viral yields recovered from A549 transfections precluded the production of usable viral stocks. We therefore proceeded with the double mutant stocks produced in Vero cells (N24A-D31N and N24D-D31H/N) for all subsequent experiments.

**Figure 2.**
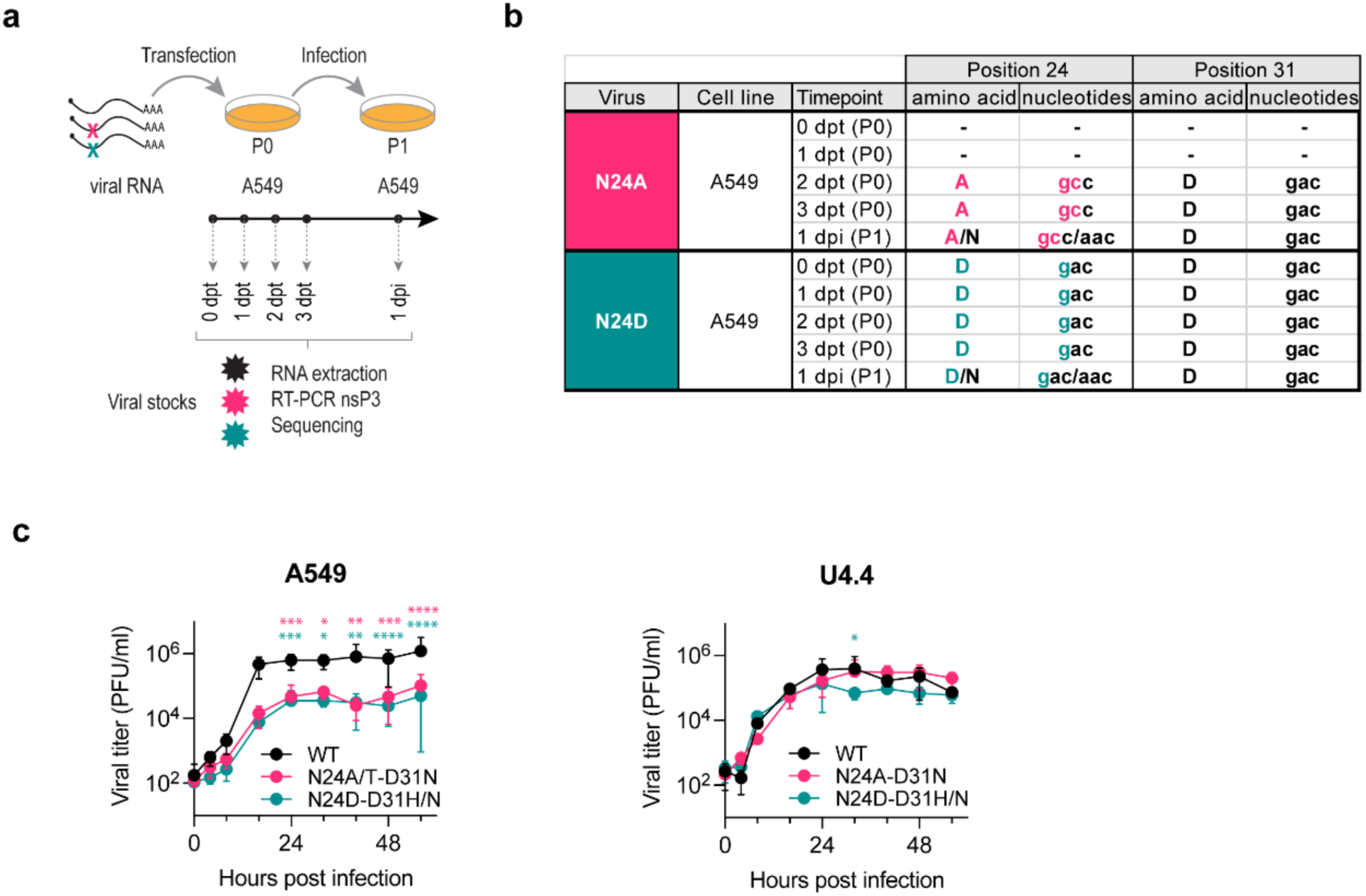
N24 catalytic mutations impair CHIKV replication in interferon-competent human cells but not in mosquito cells. a, Schematic of the experimental design. *In vitro* transcribed N24A and N24D viral RNAs were transfected into A549 cells. Cell culture supernatants were collected at 0, 1, 2, and 3 days post-transfection (P0) and at 1 day post-infection (P1) and subjected to RNA extraction, RT-PCR amplification of the region encoding the first 119 amino acids of the nsP3 macrodomain, and Sanger sequencing. **b,** Amino acid and nucleotide identities at residues 24 and 31 in N24A and N24D viruses following transfection into A549 cells. Dashes indicate timepoints where insufficient viral RNA was recovered for sequencing. Double peaks indicating coexistence of multiple nucleotides at the same position are denoted by ‘/’. Amino acid abbreviations: N, asparagine; A, alanine; D, aspartic acid. **c,** Growth kinetics of WT, N24A-D31N, and N24D-D31H/N viruses in A549 and U4.4 cells. In A549 cells, the N24A-D31N virus is labelled N24A/T-D31N to reflect the mixed alanine/threonine population detected at position 24 at 48 hours post-infection (Supplementary Fig. 1c). Cells were infected at a multiplicity of infection of 0.05. Data are plotted as mean ± SD for three independent biological experiments. Data were analyzed using a mixed-effects model with Geisser-Greenhouse correction followed by Tukey’s multiple comparison test. Pink and teal asterisks indicate significant differences compared to WT. *p < 0.05; **p < 0.01; ***p < 0.001; ****p < 0.0001.

To assess the replication capacity of the double mutant viruses in mammalian and mosquito cells, we performed growth curve experiments in A549 and U4.4 cells using these Vero-derived stocks (Fig. 2c). In A549 cells, both N24A-D31N and N24D-D31H/N mutant viruses reached significantly lower viral titers than WT at multiple timepoints across the growth curve, demonstrating that loss of macrodomain catalytic activity impairs CHIKV replication in interferon-competent human cells. In contrast, in U4.4 mosquito cells, both mutant viruses replicated to levels broadly comparably to WT across the time course, except for a significantly lower titer observed for N24D-D31H/N at 32 hours post-infection. The genetic stability of the introduced mutations was assessed by Sanger sequencing of the first 119 amino acids of nsP3 at 48 hours post-infection in both cell lines (Supplementary Fig. 1c). Both mutations remained stable, except for the Vero-derived N24A-D31N virus in A549 cells, where a mixed population of alanine and threonine was detected at position 24 (GCC/ACC). This indicates that the N24A substitution is under selective pressure in interferon-competent human cells, consistent with the partial reversion observed after transfection in the same cell line. No such change was detected in U4.4 mosquito cells.

Next, we examined the impact of the N24-D31 mutations in the context of the Indian Ocean CHIKV strain (Supplementary Fig. 1d). In human A549 cells, both mutant viruses reached significantly lower viral titers than WT at 40 hours post-infection (14-fold and 43-fold lower for N24A-D31N and N24D-D31N, respectively), and N24D-D31N also showed reduced titers at 24 hours post-infection (27-fold lower than WT). In contrast, in U4.4 mosquito cells, only N24D-D31N showed modestly reduced titers compared to WT, at 16 and 48 hours post-infection (3-fold and 5-fold lower, respectively). As observed for CHIKV-Caribbean, the presence of both N24 and D31 mutations in growth curve samples (collected at 72 hours post-infection) was confirmed by nsP3 RT-PCR and Sanger sequencing, with partial reversion at position 31 detected for N24A-D31N in U4.4 cells (Supplementary Fig. 1e).

Overall, these results demonstrate a host-dependent difference in the requirement for macrodomain catalytic activity that is independent of the viral lineage: while it is important for viral replication in interferon-competent mammalian cells, it is largely dispensable in mosquito cells. This raised the question of whether macrodomain catalytic activity is similarly dispensable during CHIKV infection of *Aedes* mosquitoes *in vivo*, and whether the selective pressure driving D31 second-site mutation emergence operates in the mosquito host.

### Loss of macrodomain catalytic activity modulates CHIKV dissemination and transmission potential in *Aedes* mosquitoes

To investigate the role of macrodomain catalytic activity in CHIKV infection of the mosquito host, we exposed laboratory colonies of *Ae. albopictus* and *Ae. aegypti* to an infectious blood meal containing Caribbean WT, N24A-D31N, or N24D-D31H/N viruses. At 2, 5, and 7 days post-infection, individual mosquitoes were dissected into bodies (to study viral dissemination) and heads (to analyze transmission potential). Prevalence of infection and viral titers were determined separately for bodies and heads at each timepoint (Fig. 3a). To assess the genetic stability of the introduced mutations and the selection pressure during mosquito infection, viral RNA from individual infected mosquitoes was extracted and the first 119 amino acids of the nsP3 gene was sequenced (Supplementary Table 2).

**Figure 3.**
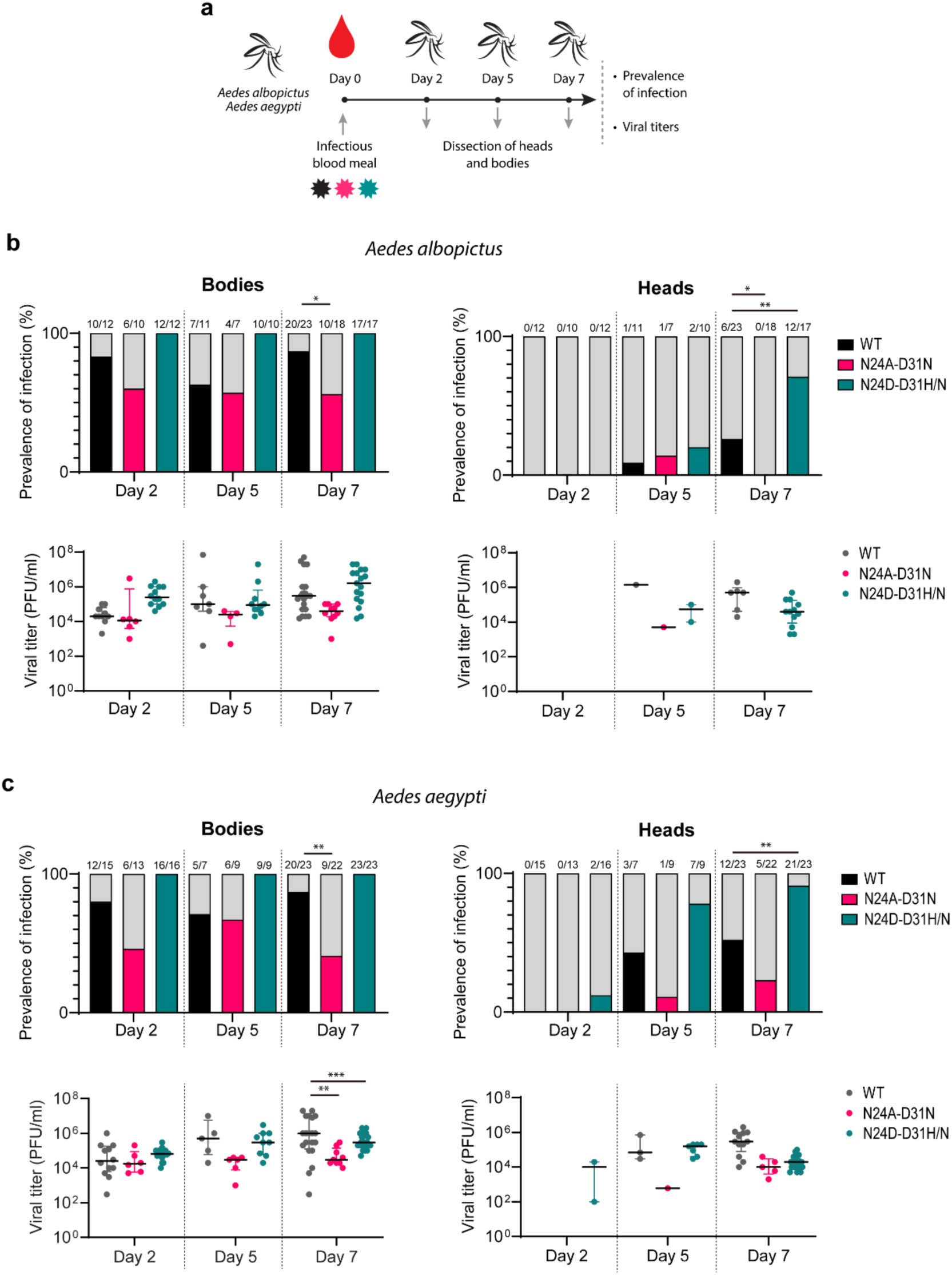
Macrodomain catalytic mutations modulate CHIKV dissemination and transmission potential in *Aedes* mosquitoes. a, Schematic of the experimental design. Laboratory colonies of *Ae. albopictus* and *Ae. aegypti* were exposed to an infectious blood meal containing WT, N24A-D31N, or N24D-D31H/N viruses. At 2, 5, and 7 days post-infection, individual mosquitoes were dissected into bodies and heads. Prevalence of infection and viral titers were determined by plaque assay. **b,** Prevalence of infection (top) and viral titers (bottom) in bodies and heads of *Ae. albopictus* mosquitoes. **c,** Prevalence of infection (top) and viral titers (bottom) in bodies and heads of *Ae. aegypti* mosquitoes. Numbers above prevalence bars indicate the number of infected individuals over the total number of mosquitoes tested. Horizontal bars represent the median and interquartile range. Individual data points represent single mosquitoes. Prevalence of infection was compared using Fisher’s exact test. Viral titers were analyzed using a two-way ANOVA followed by Tukey’s multiple comparison test. Asterisks indicate statistically significant differences compared to WT. *p < 0.05; **p < 0.01; ***p < 0.001.

The N24A-D31N mutant showed reduced dissemination and transmission potential compared to WT virus in both mosquito species. In *Ae. albopictus*, prevalence of infection in bodies was significantly lower for N24A-D31N than for WT at day 7 (56% vs 87%), and no infected heads were detected at this timepoint, compared to 26% for WT (Fig. 3b). In *Ae. aegypti*, viral titers in bodies were significantly lower for N24A-D31N than for WT at day 7, and prevalence of infection in bodies was also significantly reduced at day 7 (41% vs 87%) (Fig. 3c). Head prevalence of infection at day 7 was lower for N24A-D31N than for WT (23% vs 52%), though this difference did not reach statistical significance. Sequencing of the first 119 amino acids of nsP3 from individual infected mosquitoes confirmed that both N24A and D31N mutations remained stable throughout the infection time course in both species, with the exception of a single *Ae. aegypti* individual at day 5, in which a partial reversion towards the WT asparagine was detected at position 24, accompanied by complete reversion to the parental aspartic acid at position 31 (Supplementary Table 2).

In contrast, the N24D-D31H/N mutant showed enhanced dissemination and transmission potential compared to WT. In *Ae. albopictus*, N24D-D31H/N reached 100% prevalence of infection in bodies at all three timepoints, compared to 83%, 64%, and 87% for WT at days 2, 5, and 7 respectively. Head prevalence of infection at day 7 was significantly higher for N24D-D31H/N than for WT (71% vs 26%), while viral titers in infected heads were comparable between the two viruses (Fig. 3b). In *Ae. aegypti*, N24D-D31H/N again reached 100% prevalence of infection in bodies at all timepoints. Head prevalence of infection at day 7 was significantly higher for N24D-D31H/N than for WT (91% vs 52%), while viral titers in infected heads were comparable. Notably, despite the enhanced dissemination of N24D-D31H/N, viral titers in bodies were significantly lower for this mutant compared to WT at day 7 in *Ae. aegypti*, suggesting that enhanced dissemination is not driven by increased viral replication (Fig. 3c). Sequencing of nsP3 from individual infected mosquitoes revealed that the N24D mutation remained stable throughout the infection time course, except for a single *Ae. aegypti* individual at day 2, which showed a partial reversion toward the WT sequence at position 24 together with reversion to aspartic acid at position 31. At position 31, the viral population evolved from a predominantly D31H/N mixture at day 2 toward predominantly D31H by day 7 in both species, indicating that the D31H variant is preferentially selected in the mosquito host (Supplementary Table 2).

To determine whether the enhanced dissemination and transmission potential of the N24D-D31H/N double mutant could be attributed to the D31 mutations independently of N24 catalytic inactivation, we generated single D31H and D31N mutant viruses and assessed their replication in A549 and U4.4 cells and in *Aedes* mosquitoes (Supplementary Fig. 2). In cell culture, D31H and D31N single mutants replicated to levels broadly comparably to WT in both A549 and U4.4 cells. In A549 cells, D31H reached significantly higher titers than WT at the last timepoint, whereas in U4.4 cells both mutants showed significantly lower titers than WT at early timepoints before reaching WT levels. In both mosquito species, D31H and D31N single mutants showed no significant differences in prevalence of infection compared to WT at any timepoint, in either bodies or heads. Viral titers were also comparable to WT, with the exception of *Ae. albopictus* at day 5, where D31H reached significantly higher titers than WT in bodies and heads. The single D31 mutants did not reproduce the phenotype of the N24D-D31H/N double mutant, which was characterized by an increased prevalence of infection in bodies and heads rather than by increased viral titers. The genetic stability of the introduced D31 mutations was confirmed by Sanger sequencing of the first 119 amino acids of nsP3 from individual infected mosquitoes (Supplementary Table 3). These results demonstrate that D31 mutations alone are not sufficient to reproduce the enhanced dissemination and transmission potential of the N24D-D31H/N double mutant, suggesting that this phenotype depends on the loss of macrodomain catalytic activity at position N24.

To evaluate whether the *in vivo* effects of macrodomain mutations depend on the CHIKV strain, we infected both mosquito species with the Indian Ocean WT and mutant viruses (Supplementary Fig. 3). In *Ae. albopictus*, significant differences were observed at day 5 post-infection, with the N24D-D31N mutant reaching higher viral titers in bodies and higher prevalence of infection in heads compared to WT (Supplementary Fig. 3a). In *Ae. aegypti*, no differences in viral titers or prevalence of infection were found between WT and mutant viruses (Supplementary Fig. 3b). Of note, experimental infection of *Ae. aegypti* with Indian Ocean viruses was highly efficient, with 100% of bodies infected at all timepoints tested, which likely masked potential differences between WT and mutant viruses. Sequencing of the first 119 amino acids of nsP3 from individual infected mosquitoes confirmed that the introduced N24 mutations remained stable throughout the infection time course in both species (Supplementary Table 2). At position 31, however, the viral populations were more variable, with partial reversion toward the parental aspartic acid detected in a subset of individuals of both species. The D31H variant emerged in three *Ae. albopictus* individuals infected with the N24D-D31N virus at day 5 post-infection, mirroring the preferential selection of D31H observed in mosquitoes infected with the Caribbean strain (Supplementary Table 2).

Taken together, these results demonstrate that macrodomain catalytic mutations modulate CHIKV dissemination and transmission potential in a mutation-specific manner that is consistent across both *Aedes* species. The distinct effects are driven primarily by differences in prevalence of infection rather than viral titers in infected individuals, suggesting that macrodomain catalytic activity influences the ability of CHIKV to establish infection and disseminate within the mosquito rather than its replication level once established. The enhanced presence in heads of the N24D-D31H/N mutant in both mosquito species represents a gain-of-function phenotype. Given this finding, and in line with institutional guidance, we did not repeat experiments with this viral variant that was discarded.

### The CHIKV macrodomain catalytic activity is abolished by N24 mutations and is not rescued by D31 second-site mutations

Mosquito experiments showed that the N24D-D31H/N mutant, despite carrying a mutation at the catalytic site, enhanced CHIKV dissemination. This raised the question of whether the D31 second-site mutations might rescue catalytic activity, thereby explaining the observed phenotype. To address this, we investigated the structural and functional contributions of N24 and D31 to macrodomain catalytic activity and ADP-ribose binding capacity *in vitro*. We constructed expression plasmids encoding WT and mutant versions of the CHIKV macrodomain and purified the following recombinant macrodomain variants from bacteria: WT, N24A, N24D, D31H, D31N, N24A-D31H, N24A-D31N, N24D-D31H, and N24D-D31N (Supplementary Fig. 4a). First, we assessed the catalytic activity of recombinant macrodomains using an AMP-Glo luciferase assay^4^, in which ADP-ribose released from a MARylated protein substrate is converted to AMP and detected as a luminescent signal proportional to hydrolase activity (Fig. 4a). All N24 mutants (in either single or double combinations with D31) were catalytically inactive, exhibiting equivalent luciferase activity to the non-enzyme control. In contrast, single D31 mutants did not significantly alter catalytic activity compared to WT, demonstrating that the D31 second-site mutations do not rescue catalytic activity and therefore cannot account for the enhanced mosquito infectivity observed for the N24D-D31H/N mutant.

**Figure 4.**
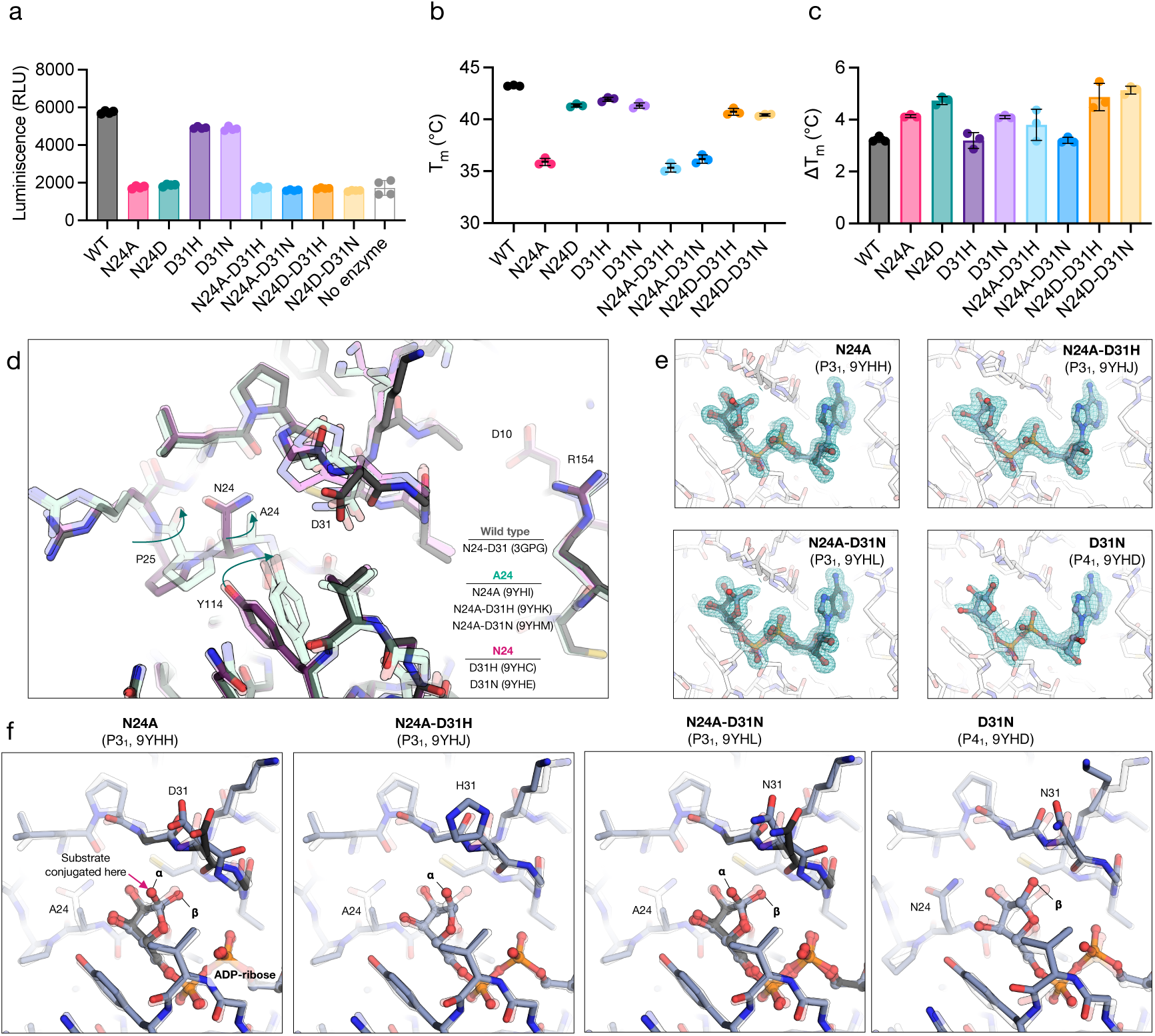
Functional and structural characterization of CHIKV nsP3 macrodomain mutants. **a**, Assays of the ADP-ribosylhydrolase activity of recombinantly expressed wild-type (WT) and mutant macrodomains. Macrodomains (200 nM) were incubated with auto-MARylated human PARP10 for 1 hour at room temperature and the production of ADP-ribose was measured using NUDT5 and an AMP-Glo luciferase assay^4^. Data are plotted mean ± SD for four technical replicates. **b**, Thermostability of CHIKV nsP3 macrodomain mutants measured by DSF using SYPRO orange. Data are plotted mean ± SD for three technical replicates. **c**, Change in CHIKV nsP3 macrodomain thermostability upon incubation with 1 mM ADP-ribose. **d,** Alignment of the crystallographic structures of WT, N24A, N24A-D31N, N24A-D31H, D31H and D31N reveals a peptide flip in P25 and a coupled shift in Y114 in structures containing the N24A mutation. For clarity, only chain A is shown (see Supplementary Fig. 5a for all chains). **e**, Difference electron density maps (F_O_-F_C_, contoured at 3 σ) calculated prior to modeling ADP-ribose. Maps for all chains are shown in Supplementary Fig. 6. **f**, Crystal structures of N24A, D31N, N24A-D31H and N24A-D31N bound to ADP-ribose. Chain A is shown for the P3_1_ structures and chain D for the P4_1_ structure. Although the ADP-ribose binding pose is conserved, there is a rotameric shift at residue 31 that accompanies ADP-ribose binding, and mutations at position 31 change the character of the exit path of the substrate suggesting a role for substrate-specific recognition.

Next, we determined the thermostability and ADP-ribose binding capacity of the mutants using differential scanning fluorimetry (DSF), which tracks thermal unfolding via a dye that increases fluorescence on binding newly exposed hydrophobic surfaces, yielding a melting temperature (Tm). The apparent Tm generally increases when a ligand binds to the protein as additional energy is required to dissociate the ligand prior to unfolding (Fig. 4b, 4c, Supplementary Table 4, Supplementary Fig. 4b). This analysis revealed that the N24A mutant is less stable (ΔTm relative to WT = -7.3°C) than the N24D mutant (ΔTm relative to WT = -1.9°C), similar to results with the SARS-CoV-2 macrodomain^4^. Both mutants also retained ADP-ribose binding capacity, generating a similar Tm shift in the presence of 1 mM ADP-ribose to WT (3.2°C for WT, 4.1°C for N24A, 4.7°C for N24D) (Fig. 4c, Supplementary Table 4). We note that DSF measures binding capacity at a fixed ligand concentration rather than binding affinity, and we cannot exclude subtle quantitative differences in ADP-ribose affinity among the mutants. The compensatory mutations had minimal effects on both stability and ADP-ribose binding capacity. The D31 mutations were neutral in the WT background and did not rescue stability in the N24-mutant backgrounds. This result suggests that although N24A is destabilizing, the compensatory mutations do not act to restore protein stability.

To examine how these mutations might alter structure and function, we determined X-ray structures of N24A, N24A-D31H, N24A-D31N, D31H and D31N (Fig. 4d, Supplementary Table 5). The structures reveal a shift in residues 24-25 in the N24A mutants, with a flip in the P25 carbonyl and a correlated shift in nearby Y114 side chain (Fig. 4d, Supplementary Fig. 5a, 5b). Given the lack of change in either catalytic or ADP-ribose binding capacity relative to the N24 mutations alone, we expected that the compensatory residues would not alter the interactions with ADP-ribose. To test this, we soaked ADP-ribose into crystals and determined structures of N24A, N24A-D31H, N24A-D31N and D31N (Fig. 4e, Supplementary Table 5, Supplementary Fig. 6). The structures show that the D31 mutants undergo a similar side chain rotation when ADP-ribose binds (Fig. 4f). We observed both the α and β anomers of ADP-ribose bound (Fig. 4f, Supplementary Fig. 5c-f): the α anomer is consistent with the substrate bound state, while both anomers are consistent with product bound^44^. The D31 mutations line the “exit” path of the residue modified with ADP-ribose, leaving them poised to influence substrate binding to the macrodomain (Fig. 4f). Collectively, these results confirm that the catalytic activity of the macrodomain is abolished by N24 mutations and is not rescued by the D31 second-site mutations, while ADP-ribose binding capacity is maintained. The structural location of the independently acquired D31 mutations at the substrate exit path raises the possibility that they influence ADP-ribose substrate recognition, potentially tuning the reader activity of the macrodomain toward specific substrates. These findings confirm that the viral phenotypes observed in mosquitoes are not a consequence of restored catalytic activity but arise in the context of a catalytically inactive macrodomain that retains ADP-ribose binding capacity.

## Discussion

Here we investigated the role of macrodomain catalytic activity in CHIKV replication and transmission between vertebrate and invertebrate hosts. While previous studies have established the importance of the alphavirus macrodomain for replication and virulence in mammalian hosts, its specific role in the mosquito vector remained largely unexplored. Our data demonstrate that macrodomain mutations directly impact viral infectivity and dissemination in *Aedes* mosquitoes *in vivo*. This reveals a previously unknown host-specific function of the macrodomain. By introducing N24A and N24D mutations into the CHIKV nsP3 macrodomain, we abolished ADP-ribosylhydrolase activity while preserving ADP-ribose binding capacity. The consistent emergence of D31 mutations specifically in CHIKV-N24 catalytic mutants, together with the selection of a second-site mutation at the equivalent residue in a Sindbis virus macrodomain double mutant^43^, suggests that residue 31 is a recurrent site of genetic compensation in alphavirus macrodomains. However, the specific selective pressures driving D31 emergence -whether catalytic compensation, protein stability effects, or other factors -remain to be definitively established. The structural location of the D31 mutations at the substrate exit path, combined with their progressive selection toward D31H in the mosquito host, raises the possibility that these mutations tune ADP-ribose substrate recognition in a host-specific manner.

The observation that N24 mutations are poorly tolerated in interferon-competent human cells but not in mosquito cells is consistent with the established role of macrodomain catalytic activity in counteracting PARP-mediated antiviral responses in mammals and promoting viral virulence *in vivo*^9,12,45^. Of note, although both CHIKV lineages replicated productively in A549 cells under our experimental conditions, infection did not result in detectable cytopathic effects, consistent with previous reports describing a limited permissiveness of this cell line to CHIKV^46–48^. This relatively restrictive context may in fact favor the detection of replication differences driven by the interferon response, which is central to macrodomain function in mammalian cells. Our observations are also in line with the recurrent loss of N24 in mosquito-specific alphaviruses^35,36^, which replicate exclusively in an environment lacking interferon-driven PARP responses. Our experimental data extend these observations by directly demonstrating, for the first time *in vivo*, that catalytic inactivation of the macrodomain modulates CHIKV dissemination and transmission potential in *Aedes* mosquitoes in a mutation-specific manner. While the N24A-D31N mutation impaired viral dissemination, the N24D-D31H/N mutation enhanced infectivity and transmission potential in both *Ae. albopictus* and *Ae. aegypti*. Importantly, single D31 mutant viruses did not recapitulate this gain-of-function phenotype, suggesting that the enhanced dissemination and transmission potential phenotype depends on loss of catalytic activity at position N24 rather than the D31 changes alone. We note, however, that the full viral genomes were not sequenced, so additional mutations outside the macrodomain region cannot be excluded. This differential evolutionary trajectory provides insights into how dual-host arboviruses balance conflicting host immune pressures.

Our results can be compared with earlier studies of alphavirus macrodomain mutants. In CHIKV 181/25, mutations that abolished hydrolase activity (D10A, G32E and G112E) did not yield viable virus and reverted to the parental residue^12^. Reversion of G32E occurred in both mammalian BHK-21 and mosquito C6/36 cells, leading the authors to conclude that hydrolase activity is required in both hosts^12^. Our data are consistent with this at the level of selection: although single N24A and N24D genomes were detected shortly after transfection, they could not be propagated into usable viral stocks in mammalian cells. In our system, both partial reversion at position 24 and second-site mutations at residue 31 were observed, often within the same population. Importantly, the viruses that could be propagated as stocks retained the N24 substitution together with a mutation at residue 31, and the Caribbean double mutants replicated to levels comparable to WT in U4.4 mosquito cells despite having no detectable catalytic activity. The equivalent Indian Ocean mutants showed only modest reductions in the same cells. This indicates that catalytic activity itself is largely dispensable in mosquito cells, provided the macrodomain is compensated at residue 31.

The nature of the compensation differs between viruses. In Sindbis virus, single mutation of the equivalent asparagine 24 is well tolerated^11,43^, and the second-site change at residue 31 was selected only when this mutation was combined with a second substitution in the ADP-ribose binding site^43^. That change, from glutamate to glycine, made the macrodomain more similar to other macrodomains, in which glycine is the common residue at this position^43^. In CHIKV, by contrast, a single N24 substitution was already sufficient to select mutations at residue 31, and the selected residues were asparagine or histidine, neither of which is common at this position in viral macrodomains (Fig. 1e, 1f). The compensation observed here therefore appears specific to the CHIKV macrodomain rather than a general return toward an ancestral sequence.

The gain-of-function enhancement of mosquito transmission potential observed for N24D-D31H/N is unexpected and its molecular mechanism remains to be determined. The finding that viral titers in infected mosquito bodies were lower for N24D-D31H/N than for WT despite higher prevalence of infection suggests that this mutant is more efficient at establishing infection across midgut cells rather than replicating to higher levels once infection is established. One possibility is that the catalytically inactive but ADP-ribose binding-competent macrodomain interacts differently with mosquito host proteins, potentially influencing the higher-order structures formed by nsP3 that are needed to interact with host factors^49^ or organize tubular structures containing viral genomic RNA^50^. The progressive selection of D31H over D31N in the mosquito host further suggests that subtle differences in ADP-ribose substrate recognition may influence viral fitness in the mosquito environment in ways that are not yet understood. Notably, which proteins are ADP-ribosylated under interferon signaling in human cells is only just emerging^51^, and the potential interactome in mosquito cells has not been established. This knowledge gap highlights the challenge in understanding how the D31 mutations redirect macrodomain specificity in a host-dependent manner.

Our findings have important implications for the development of macrodomain-targeted antivirals. While macrodomain inhibitors targeting catalytic activity are promising therapeutic candidates for mammalian infections^14^, our results demonstrate that the function of this domain is fundamentally host-dependent. The unexpected gain-of-function enhancement of virus dissemination and transmission potential in mosquitoes by a catalytically inactive macrodomain variant underscores the importance of evaluating antiviral strategies across the full host range of dual-host viruses. Compounds that inhibit macrodomain catalytic activity may effectively suppress viral replication in mammals while having unpredictable effects on transmission by the mosquito vector. A deeper understanding of macrodomain function across hosts is therefore essential for the rational design of broad-spectrum antivirals targeting this domain.

## Methods

### Construction of recombinant viruses

To introduce the N24A and N24D mutations into the nsP3 gene of the CHIKV-Caribbean^52^ and -Indian Ocean^53^, 5’-phosphorylated primers were used to amplify the full-length plasmid template via inverse PCR using Q5 High Fidelity DNA Polymerase (New England Biolabs). Forward primers contained 1-2 mismatches relative to the WT CHIKV sequences to incorporate the desired nucleotide changes into the PCR product. N24A mutations were introduced using the forward primer 5’-GCCCCTCGCGGGTTACCG-3’. N24D mutations were introduced using the forward primer 5’-GACCCTCGCGGGTTACCG-3’. The reverse primer for both mutants was 5’-GGCGGCGTTGACCACGCA-3’. To introduce the D31H and D31N mutations, the same inverse PCR strategy was used. The forward primer 5’-ACCCGGTAACCCGCGAGGG-3’ was used for both mutants. The reverse primer for D31H was 5’-CACGGTGTTTGCAAGGCAG-3’ and for D31N was 5’-AACGGTGTTTGCAAGGCAG-3’. All primers were 5’-phosphorylated. Purified PCR products were blunt-end ligated with T4 DNA ligase and transformed into competent *E. coli* cells. Colonies were miniprepped, and plasmid sequences were verified by whole-plasmid sequencing (Eurofins).

The DNAs of the recombinant constructs were linearized by digestion with NotI enzyme and used as templates for transcription by SP6 polymerase in the presence of GpppG cap structure analog, using the mMessage mMachine transcription kit (Thermo Fisher). The RNAs were gel-quantified and used for transfection in cell culture.

### Cells, viral transfections and infections

Mammalian cell lines (Vero, A549 and BHK-21) were maintained in Dulbecco’s Modified Eagle Medium (DMEM) supplemented with 10% fetal bovine serum (FBS) and 1% penicillin/streptomycin (P/S) at 37°C with 5% CO_2_. *Ae. albopictus* cell lines (C6/36 and U4.4) were maintained in Leibovitz L-15 medium supplemented with 10% FBS, 1% P/S, 1% non-essential amino acids, and 1% tryptose phosphate broth at 28°C with 5% CO_2_.

For virus stock production, Vero cells were grown to 60-70% confluence in T75 flasks and transfected with 10 μg of in vitro transcribed viral RNAs using Lipofectamine 2000 (Invitrogen). Cell culture supernatants were collected at one day post-transfection and used to infect fresh Vero cells. Viral stocks were harvested from cell culture supernatants at one and two days post-infection and quantified by plaque assay.

### Plaque assays

Viruses from culture supernatants were quantified by plaque assays to determine viral yields. Vero cells (1 × 10^5^ per well) were seeded into 24-well plates and allowed to attach overnight. Viral stocks were serially diluted, and 0.1 mL was added to the cells, followed by a 1-hour incubation. An overlay medium (1× DMEM, 2% FBS, 1% P/S, and 0.8% agarose) was then added to each well. Cells were fixed three days post-infection with 4% formaldehyde and stained with crystal violet for plaque visualization.

### Assessment of mutations in viral macrodomains

To assess the genetic stability of N24A and N24D mutations in the different cell lines tested (Vero, BHK, A549 and C6/36), cells were transfected in T25 flasks with 3 μg of *in vitro* transcribed viral RNAs. Cell culture supernatants were harvested at 0, 1, 2, and 3 days post-transfection (P0) for total RNA extraction, nsP3 RT-PCR amplification, and Sanger sequencing. Viruses recovered from supernatants were used to re-infect fresh A549 cells, and cell culture supernatants from this passage (P1) were also harvested for total RNA extraction, nsP3 RT-PCR, and sequencing.

Viral RNAs were extracted from culture supernatants using TRIzol (Invitrogen) and used for reverse transcription (RT) reactions with oligo-dT primers (Maxima H Minus RT, Thermo Fisher Scientific). PCR reactions were performed using the forward primer 5’-GCAGTTTTGACAATGGCAGAAG-3’ and the reverse primer 5’-TGTCTTTCCCTCCTGAGTATACAC-3’ (DreamTaq DNA Polymerase, Thermo Fisher Scientific). The size and quality of the amplicons were assessed by electrophoresis on a 1.5% agarose gel. The 453-bp amplicons were cleaned up using ExoSAP-IT™ PCR Product Cleanup Reagent (Thermo Fisher Scientific) and submitted for Sanger sequencing (Eurofins).

For each PCR sample, individual Sanger chromatograms were visually inspected to assess sequence quality and detect double peaks. All high-quality sequences were aligned against the WT CHIKV nsP3 gene to identify reversions of introduced mutations and detect new mutations.

### Sequence alignment and phylogram construction

Protein sequences of the nsP3 macrodomain from 14 alphaviruses and 3 other human viruses were downloaded from GenBank and the Protein Data Bank (PDB). The protein sequence identifiers and full names of the viruses are detailed in Supplementary Table 1. Multiple sequence alignment and phylogenetic tree construction were performed using the online Clustal Omega tools^54^.

### Growth curves

Sub-confluent A549 and U4.4 cells in six-well plates were infected with equal amounts of WT or mutant CHIKVs recovered from Vero cells. Infections were performed using a multiplicity of infection (MOI) of 0.05 in 500 μL of phosphate-buffered saline (PBS). One hour post-infection, cells were washed three times with PBS, and 2 mL of growth media were added. At predetermined time points, cell culture supernatants were collected and stored at -80°C. Virus quantification was performed by plaque assay on Vero cells using serial dilutions of the supernatants. Growth curves were performed in three independent biological experiments for Caribbean double mutant viruses and two independent biological experiments for Caribbean single D31 mutant viruses and Indian Ocean double mutants. To confirm genetic stability of the introduced mutations, cell supernatants from the 48 hour timepoint were extracted using TRIzol, amplified by RT-PCR, and analyzed by Sanger sequencing as previously described.

### Mosquito rearing

Laboratory colonies of *Ae. aegypti* mosquitoes (17th generation; collected originally in Kamphaeng Phet Province, Thailand) and *Ae. albopictus* (19th generation; collected originally in Phu Hoa, Binh Duong Province, Vietnam) were used. The insectary conditions for mosquito maintenance were 28°C, 70% relative humidity, and a 12-h-light and 12-h-dark cycle. Adults were maintained with permanent access to a 10% sucrose solution. Adult females were offered commercial rabbit blood (BCL, Boisset-Saint-Priest, France) twice a week through a membrane feeding system (Hemotek Ltd.).

### Ethics

Human blood samples to prepare mosquito artificial infectious bloodmeals were supplied by healthy adult volunteers at the ICAReB biobanking platform (BB-0033-00062/ICAReB platform/Institut Pasteur, Paris/BBMRI AO203/[BIORESOURCE]) of the Institut Pasteur in the CoSImmGen and Diagmicoll protocols, which had been approved by the French Ethical Committee Ile-de-France I. The Diagmicoll protocol was declared to the French Research Ministry under reference 343 DC 2008-68 COL 1. All adult subjects provided written informed consent.

### Experimental infections of mosquitoes

Infection assays were performed with 7-to 10-day-old females starved for 24 h prior to infection in a biosafety level 3 (BSL-3) laboratory. Mosquitoes were offered the infectious blood meal for 30 min through a membrane feeding system (Hemotek Ltd.) set at 37°C with a piece of desalted pig intestine as the membrane. The blood meal was composed of washed human erythrocytes resuspended in phosphate-buffered saline mixed 2:1 with a prediluted viral stock and supplemented with 10 mM ATP (Sigma-Aldrich). The viral stock was prediluted in Leibovitz L-15 medium with 0.1% sodium bicarbonate (Gibco) to reach an infectious titer ranging from 1 × 10^6^ to 1 × 10^7^ plaque-forming units. Following the blood meal, fully engorged females were selected and incubated at 28°C with 70% relative humidity and under a 12-h-light and 12-h-dark cycle with permanent access to 10% sucrose. At different times post-infection, mosquitoes were cold anesthetized for dissection into bodies and heads. Body parts were homogenized in microtubes containing steel beads (5-mm diameter) and 300 μl of DMEM supplemented with 2% FBS using a TissueLyser II instrument (Qiagen) at 30 shakes/s for 2 min. Homogenates were clarified by centrifugation and stored at -80°C until further processing. Viral titers in individual samples were determined by plaque assay. For the assessment of mutations in the nsP3 macrodomain, RNA extracted with TRIzol from homogenates was used for RT-PCR and Sanger sequencing as previously described.

### Protein purification

Plasmid DNA (pET22b(+)) encoding His_6_-tagged wild type or mutant (N24A, N24D, D31H, D31N, N24A-D31H, N24A-D31N, N24D-D31H, N24D-D31N) macrodomains was transformed into BL21 (DE3) *Escherichia coli*. After overnight growth at 37°C (LB-agar, 100 μg/mL carbenicillin), single colonies were used to inoculate small scale cultures (10 ml LB, 100 μg/mL carbenicillin). Following overnight growth at 37°C, large scale cultures (1 L LB, 100 μg/mL carbenicillin) were inoculated and grown at 37°C until an OD600 of 0.8, at which point protein expression was induced with 0.2 mM IPTG at 18°C overnight. Cells were harvested by centrifugation (4000 g, 20 min, 4°C) and cell pellets were briefly washed with phosphate-buffered saline before being re-centrifuged (4000 g, 20 min, 4°C). The cell pellets were frozen at -80°C until further purification. Thawed cell pellets were resuspended in 25 mL Lysis buffer (50 mM Tris pH 8.0, 300mM NaCl, 20 mM imidazole, 5% glycerol, 0.1% Triton X100) and 1/2 tablet of protease inhibitor (EDTA-free cOmplete, Roche). Cells were lysed by sonication (2 min) and cell debris was collected by centrifugation (30,000 g, 30 min, 4°C). The supernatant was filtered (0.45 μm cellulose acetate) and loaded on a 5 mL HisTrap HP column pre-equilibrated with buffer A (50 mM Tris pH 8.0, 150 mM NaCl, 20mM imidazole). Bound protein was eluted via a linear gradient of buffer B (50 mM Tris pH 8.0, 150 mM NaCl, 300 mM imidazole). Eluted protein was pooled, concentrated and purified via size exclusion chromatography (HiLoad 16/600 Superdex 75, Cytiva) in 10 mM Tris pH 8.0 and 300 mM NaCl. Relevant fractions were pooled, concentrated and stored at -80°C.

### AMP-Glo assay

The ADP-ribosylhydrolase activity of CHIKV nsP3 macrodomain wild type and mutants was determined using auto-MARylated human PARP10 as a substrate following the method described previously^4^. ADP-ribose produced by the macrodomain is hydrolyzed by NUDT5 phosphodiesterase to AMP, which was detected using the commercially available AMP-Glo kit (Promega, V5011). Recombinant human PARP10 catalytic domain (residues 819–1007) and human NUDT5 (residues 1–219) were expressed in *E. coli* and purified as described previously^4^. Reactions were prepared to a final volume of 8 μl in assay buffer (20 mM HEPES pH 7.5, 100 mM NaCl, 10 mM MgCl_2_) and contained 100 nM NUDT5, 10 μM MARylated PARP10 and 200 nM CHIKV nsP3 macrodomain or buffer. The AMP concentration was quantified using a BioTek Synergy H1 plate reader after incubation of the reactions at room temperature for 1 hour. Enzyme concentration was calibrated so that no more than 20% of the substrate was consumed during the reaction. All reactions were performed in technical quadruplicates. Data were plotted using GraphPad Prism.

### Crystallography

CHIKV nsP3 macrodomain mutants were crystallized using hanging drop vapor diffusion and MRC 2-well crystallization plates. Crystals grew at room temperature from drops contained 200 nL protein and 200 nL reservoir. Crystallization conditions are reported in Supplementary Table 5. To obtain structures of mutants bound to ADP-ribose, crystals were soaked in reservoir solutions supplemented with 20 mM ADP-ribose. Crystals were vitrified in liquid nitrogen without additional cryoprotection. Data were collected at 100 K using beam line 8.3.1 of the Advanced Light Source. Data were indexed, integrated and scaled with XDS^55^ and merged with Aimless^56^. Phases were obtained by molecular replacement run with Phaser^57^ using chain A from PDB 3GPG as a search model. Initial models were improved by iterative cycles of model building using COOT^58^ and refinement with phenix.refine^59^. During initial rounds of refinement, waters were automatically added using phenix.refine. During final rounds of refinement, hydrogens were added to protein coordinates using phenix.ready_set and refined using a riding model. B-factors were refined anisotropically for non-hydrogen atoms except for the structure of the D31N mutant that crystallized in P4_1_ (PDB 9YHD). Coordinates and restraints for ADP-ribose were generated with phenix.elbow^60^. Data collection and refinement statistics are summarized in Supplementary Table 5.

### DSF

Differential scanning fluorimetry (DSF) was performed to determine the thermal stability of the target protein using SYPRO Orange dye (Thermo Fisher Scientific. The protein was diluted to a final concentration of 3 µM in assay buffer consisting of 10 mM Tris pH 8.0 and 150 mM NaCl. SYPRO Orange was used at a final concentration of 5×, prepared from a 5000x stock in DMSO according to the manufacturer’s instructions.

Each 20 µl reaction mixture was prepared in a white 96-well PCR plate by combining protein, buffer, dye and ADPr (final concentrations of 0 mM or 1 mM). Samples were mixed gently, sealed with optical film, and briefly centrifuged to eliminate bubbles. All reactions were performed in technical triplicate.

Thermal denaturation was monitored using a CFX96 Real-Time PCR Detection System (Bio- Rad). The temperature was increased from 4°C to 70°C at a rate of 1°C/min, and fluorescence was measured every minute using the FRET channel. The melting temperature (Tₘ) was determined by uploading the data to DSF World using fitting model 2^61^. Data were plotted using GraphPad Prism.

## Resource availability

### Materials availability

Plasmids generated in this study are available from the lead contact upon reasonable request.

### Data and code availability

All data reported in this paper are contained within the main text, figures, and supplementary materials. Crystallographic coordinates and structure factors have been deposited in the PDB with the following accession codes: 9YHC, 9YHD, 9YHE, 9YHF, 9YHG, 9YHH, 9YHI, 9YHJ, 9YHK, 9YHL and 9YHM.

## Acknowledgments

This work was supported by the National Institutes of Health (NIAID Antiviral Drug Discovery (AViDD) grant U19AI171110) to M-C.S., A.A. and J.S.F.; the French Government’s Investissement d’Avenir program, Laboratoire d’Excellence Integrative Biology of Emerging Infectious Diseases (grant ANR-10-LABX-62-IBEID), Fondation iXcore -iXlife -iXblue Pour La Recherche and the Explore Donation, MIE project to M-C.S. The authors thank Diego E. Alvarez for critical reading and insight into data interpretation.

## Author contributions

Conceptualization: E.S.B., J.S.F, M-C.S

Data curation: E.S.B., L.B., A.H-L, J.N., G.J.C.

Formal analysis: E.S.B., L.B., G.J.C.

Funding acquisition: A.A. J.S.F, M-C.S

Investigation: E.S.B., L.B., A.H-L, J.N., G.J.C.

Methodology: E.S.B., J.S.F

Project administration: A.A., J.S.F, M-C.S

Resources: A.A., J.S.F, M-C.S

Supervision: J.S.F, M-C.S

Validation: E.S.B., L.B., A.H-L, G.J.C., J.S.F, M-C.S

Visualization: E.S.B., G.J.C.

Writing – original draft: E.S.B.

Writing – review & editing: E.S.B, A.A., J.S.F, M-C.S

## Supplemental information

Figures S1-S6 and Tables S1-S5.

