## Supplementary material for "Loss of macrodomain catalytic activity modulates Chikungunya virus dissemination and transmission potential in *Aedes* mosquitoes"

Supplementary Figure 1

**a**

| Virus | Cell line | Timepoint | Position 24 amino acid | Position 24 nucleotides | Position 31 amino acid | Position 31 nucleotides |
|---|---|---|---|---|---|---|
| WT | BHK-21 | 0 dpt (P0) | - | - | - | - |
| | | 1 dpt (P0) | N | aac | D | gac |
| | | 2 dpt (P0) | N | aac | D | gac |
| | | 3 dpt (P0) | N | aac | D | gac |
| | | 1 dpi (P1) | N | aac | D | gac |
| | C6/36 | 0 dpt (P0) | - | - | - | - |
| | | 1 dpt (P0) | N | aac | D | gac |
| | | 2 dpt (P0) | N | aac | D | gac |
| | | 3 dpt (P0) | N | aac | D | gac |
| | | 1 dpi (P1) | N | aac | D | gac |
| N24A | BHK-21 | 0 dpt (P0) | A | gcc | D | gac |
| | | 1 dpt (P0) | A | gcc | D | gac |
| | | 2 dpt (P0) | A | gcc | D | gac |
| | | 3 dpt (P0) | A | gcc | D | gac |
| | | 1 dpi (P1) | A | gcc | D/N | gac/aac |
| | C6/36 | 0 dpt (P0) | A | gcc | D | gac |
| | | 1 dpt (P0) | A | gcc | D | gac |
| | | 2 dpt (P0) | A | gcc | D | gac |
| | | 3 dpt (P0) | - | - | - | - |
| | | 1 dpi (P1) | A | gcc | D/N | gac/aac |
| N24D | BHK-21 | 0 dpt (P0) | D | gac | D | gac |
| | | 1 dpt (P0) | D | gac | D | gac |
| | | 2 dpt (P0) | D/N | gac/aac | D/N | gac/aac |
| | | 3 dpt (P0) | D/N | gac/aac | D/N | gac/aac |
| | | 1 dpi (P1) | D/N | gac/aac | D/N | gac/aac |
| | C6/36 | 0 dpt (P0) | D | gac | D | gac |
| | | 1 dpt (P0) | D | gac | D | gac |
| | | 2 dpt (P0) | D | gac | D | gac |
| | | 3 dpt (P0) | D | gac | D | gac |
| | | 1 dpi (P1) | D | gac | D | gac |

**d**

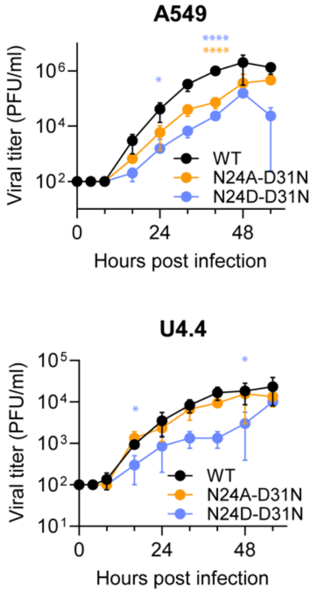


**b**

| Virus | Cell line | Timepoint | Position 24 amino acid | Position 24 nucleotides | Position 31 amino acid | Position 31 nucleotides |
|---|---|---|---|---|---|---|
| WT | BHK-21 | 1 dpt (P0) | N | aac | D | gac |
| | | 3 dpt (P0) | N | aac | D | gac |
| | | 3 dpi (P1) | N | aac | D | gac |
| | Vero | 3 dpi (P2) | N | aac | D | gac |
| N24A | BHK-21 | 1 dpt (P0) | A | gcc | D | gac |
| | | 3 dpt (P0) | A | gcc | D | gac |
| | | 3 dpi (P1) | A | gcc | D/N | gac/aac |
| | Vero | 3 dpi (P2) | A | gcc | D/N | gac/aac |
| N24D | BHK-21 | 1 dpt (P0) | D | gac | D | gac |
| | | 3 dpt (P0) | D | gac | N | aac |
| | | 3 dpi (P1) | D | gac | N | aac |
| | Vero | 3 dpi (P2) | D | gac | N | aac |

**c**

| Virus | Cell line | Timepoint | Position 24 amino acid | Position 24 nucleotides | Position 31 amino acid | Position 31 nucleotides |
|---|---|---|---|---|---|---|
| WT | A549 | 48 hpi | N | aac | D | gac |
| | U4.4 | 48 hpi | N | aac | D | gac |
| N24A-D31N | A549 | 48 hpi | A/T | gcc/acc | N | aac |
| | U4.4 | 48 hpi | A | gcc | N | aac |
| N24D-D31H/N | A549 | 48 hpi | D | gac | H/N | cac/aac |
| | U4.4 | 48 hpi | D | gac | H/N | cac/aac |

**e**

| Virus | Cell line | Timepoint | Position 24 amino acid | Position 24 nucleotides | Position 31 amino acid | Position 31 nucleotides |
|---|---|---|---|---|---|---|
| WT | A549 | 72 hpi | N | aac | D | gac |
| | U4.4 | 72 hpi | N | aac | D | gac |
| N24A-D31N | A549 | 72 hpi | A | gcc | N | aac |
| | U4.4 | 72 hpi | A | gcc | D/N | gac/aac |
| N24D-D31N | A549 | 72 hpi | D | gac | N | aac |
| | U4.4 | 72 hpi | D | gac | N | aac |

**Supplementary Figure 1. Emergence and genetic stability of macrodomain mutations across cell lines and CHIKV lineages. a,** Amino acid and nucleotide identities at residues 24 and 31 in WT,

N24A and N24D Caribbean-CHIKV (Asian lineage) viruses following transfection of viral RNAs into BHK-21 mammalian cells and C6/36 mosquito cells. Cell culture supernatants were collected at zero, one, two and three days post-transfection (dpt, P0) and at one day post-infection (dpi, P1), and subjected to RNA extraction, RT-PCR amplification of the region encoding the first 119 amino acids of the nsP3 macrodomain, and Sanger sequencing. **b,** Amino acid and nucleotide identities at residues 24 and 31 in WT, N24A and N24D Indian Ocean-CHIKV (ECSA lineage) viruses. Viral RNAs were transfected into BHK-21 cells, with supernatants collected at one and three days post-transfection (P0). Viruses were then passaged once in BHK-21 cells (P1) and a second time in Vero cells (P2), with supernatants collected at three days post-infection at each passage. **c,** Amino acid and nucleotide identities at residues 24 and 31 in WT, N24A-D31N, and N24D-D31H/N Caribbean-CHIKV viruses following infection of A549 and U4.4 cells. Supernatants were collected at 48 hours post-infection. **d,** Growth kinetics of Indian Ocean-CHIKV WT, N24A-D31N and N24D-D31N viruses in A549 and U4.4 cells. Cells were infected at a multiplicity of infection of 0.05. Data are plotted as mean ± SD. n = 3 for 0, 4, 8, 16, 32, 40 and 56 hpi, and n = 6 for 24 and 48 hpi. Data were analyzed using a mixed-effects model with Geisser-Greenhouse correction followed by Tukey's multiple comparison test. Orange and blue asterisks indicate significant differences compared to WT. *p < 0.05; ****p < 0.0001.**e,** Amino acid and nucleotide identities at residues 24 and 31 in the Indian Ocean-CHIKV viruses shown in d, recovered from A549 and U4.4 supernatants at 72 hpi. In all sequencing panels, samples were processed as described in **a**. Dashes indicate timepoints where insufficient viral RNA was recovered for sequencing. Double peaks indicating coexistence of multiple nucleotides at the same position are denoted by '/'. Amino acid abbreviations: N, asparagine; A, alanine; D, aspartic acid; H, histidine; T, threonine.

Supplementary Figure 2

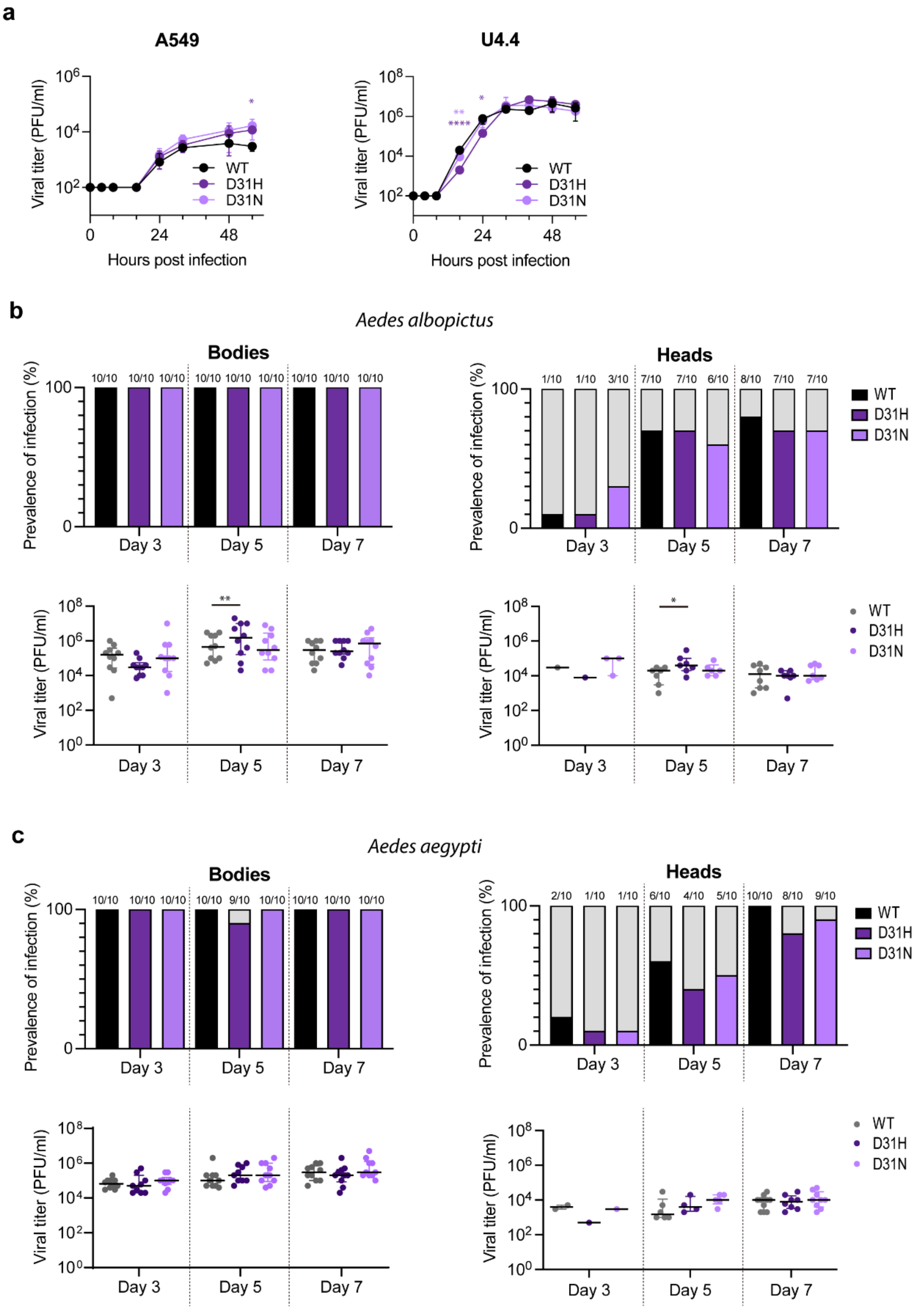


**Supplementary Figure 2. Replication and transmission potential of single D31 mutant viruses. a,** Growth kinetics of WT, D31H, and D31N viruses in A549 and U4.4 cells. Cells were infected at a multiplicity of infection of 0.05. Data are plotted as mean ± SD for two independent biological experiments. Data were analyzed using a mixed-effects model with Geisser-Greenhouse correction followed by Tukey's multiple comparison test. Purple and light purple asterisks indicate significant differences compared to WT. **b,** Prevalence of infection (top) and viral titers (bottom) in bodies and heads of *Ae. albopictus* mosquitoes infected with WT, D31H, or D31N viruses. **c,** Prevalence of infection (top) and viral titers (bottom) in bodies and heads of *Ae. aegypti* mosquitoes infected with WT, D31H, or D31N viruses. Numbers above prevalence bars indicate the number of infected individuals over the total number of mosquitoes tested. Horizontal bars represent the median and interquartile range. Individual data points represent single mosquitoes. Prevalence of infection was compared using Fisher's exact test. Viral titers were analyzed using a two-way ANOVA followed by Tukey's multiple comparison test. Asterisks indicate statistically significant differences compared to WT. *$p < 0.05$; **$p < 0.01$; ****$p < 0.0001$.

Supplementary Figure 3

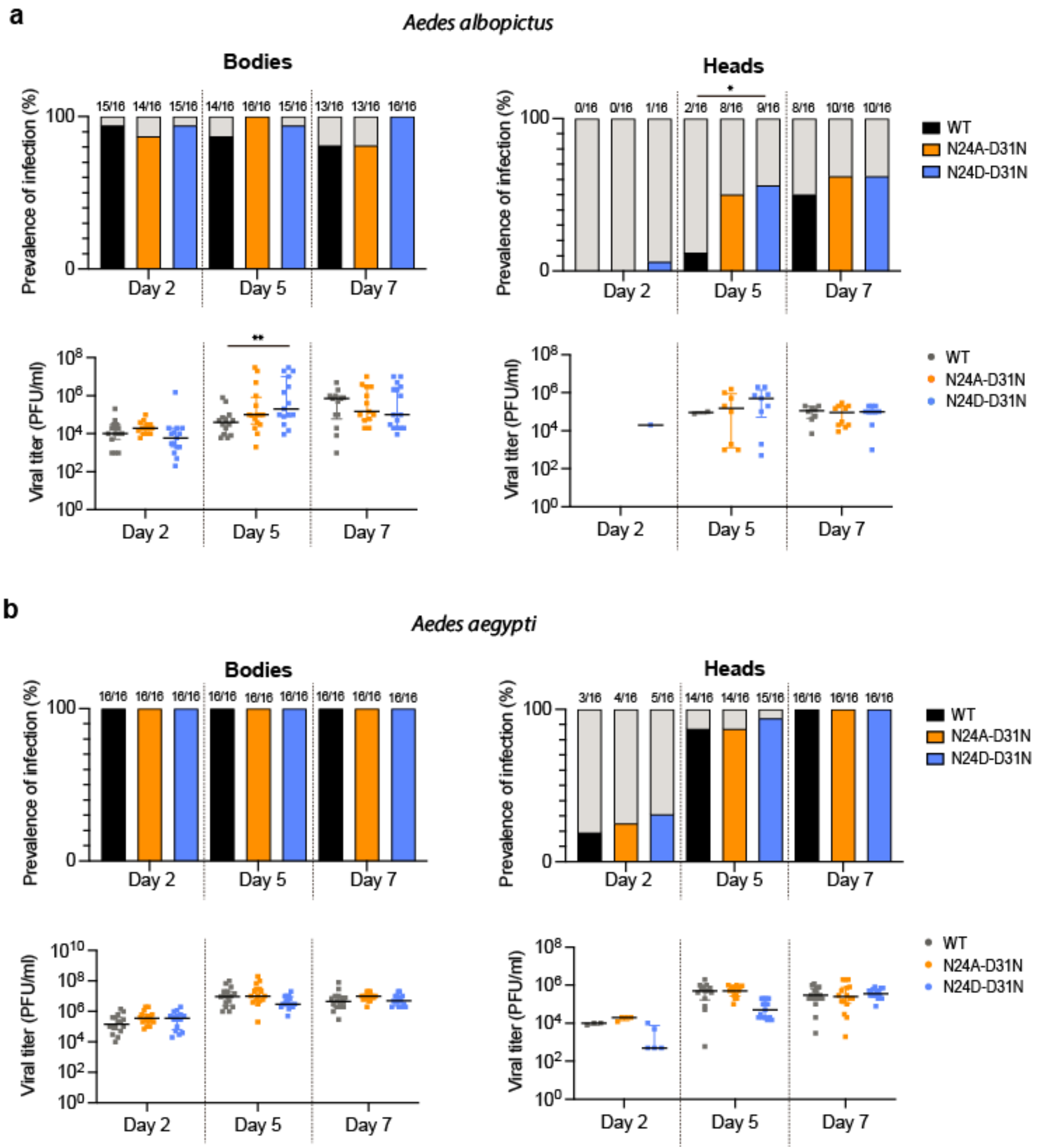


**Supplementary Figure 3. Replication and transmission potential of Indian Ocean-CHIKV macrodomain mutants in *Aedes* mosquitoes. a,** Prevalence of infection (top) and viral titers (bottom) in bodies and heads of *Ae. albopictus* mosquitoes infected with Indian Ocean WT, N24A-D31N or N24D-D31N viruses, at 2, 5 and 7 days post-infection. **b,** Prevalence of infection (top) and viral titers (bottom) in bodies and heads of Ae. aegypti mosquitoes infected with the same viruses. Numbers above prevalence bars indicate the number of infected individuals over the total number of mosquitoes tested. Horizontal bars represent the median and interquartile range. Individual data points represent single mosquitoes. Prevalence of infection was compared using Fisher's exact test. Viral titers were analyzed using a two-way ANOVA followed by Tukey's multiple comparison test. Asterisks indicate statistically significant differences compared to WT. *p < 0.05; **p < 0.01.

Supplementary Figure 4

**a**

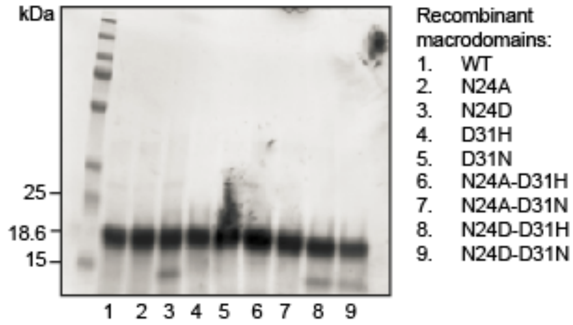


**b**

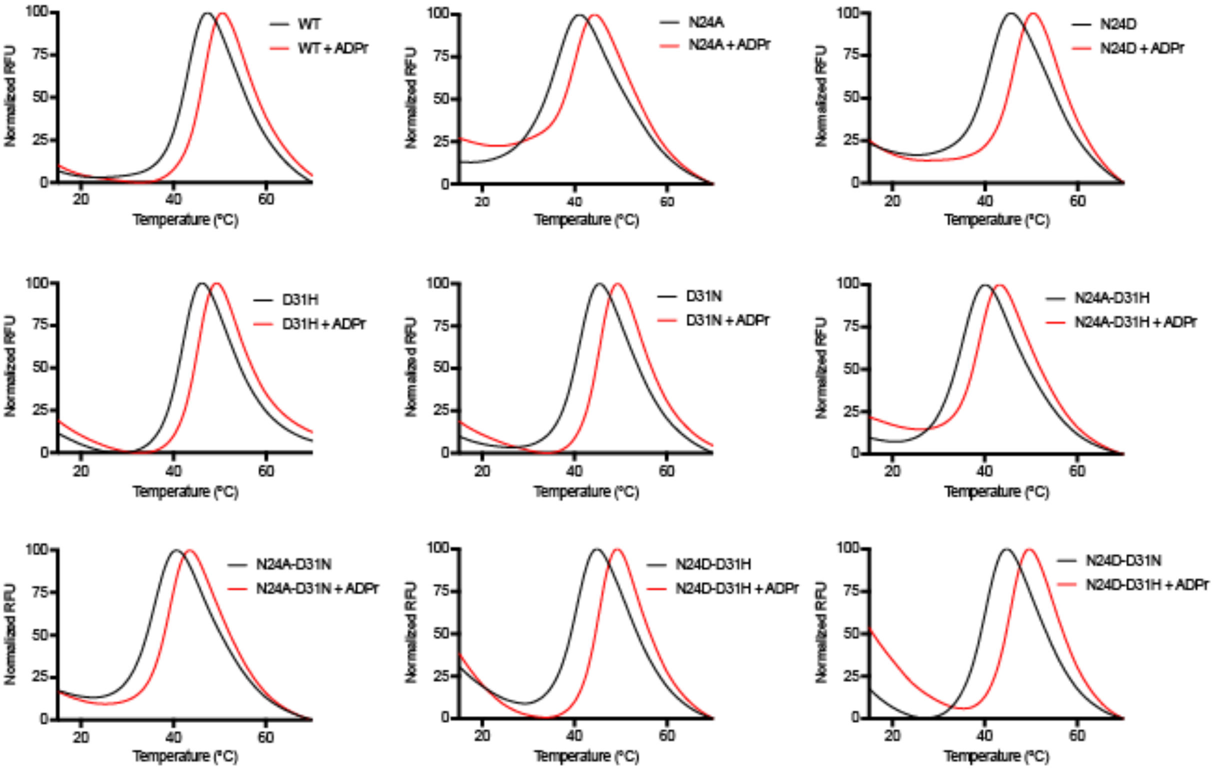


**Supplementary Figure 4. Expression of recombinant macrodomains and ADP-ribose binding assays. a**, SDS-PAGE analysis of CHIKV nsP3 macrodomain mutants recombinantly expressed and purified from *E. coli*. The expected molecular weight of the His6-tagged recombinant macrodomain is ~18.6 kDa. The minor low-molecular-weight contaminant in lanes 3, 8, and 9 is unlikely to affect the downstream results. **b**, Plot showing change in relative fluorescence upon incubation of CHIKV nsP3 macrodomain mutants with SYPRO orange.

Supplementary Figure 5

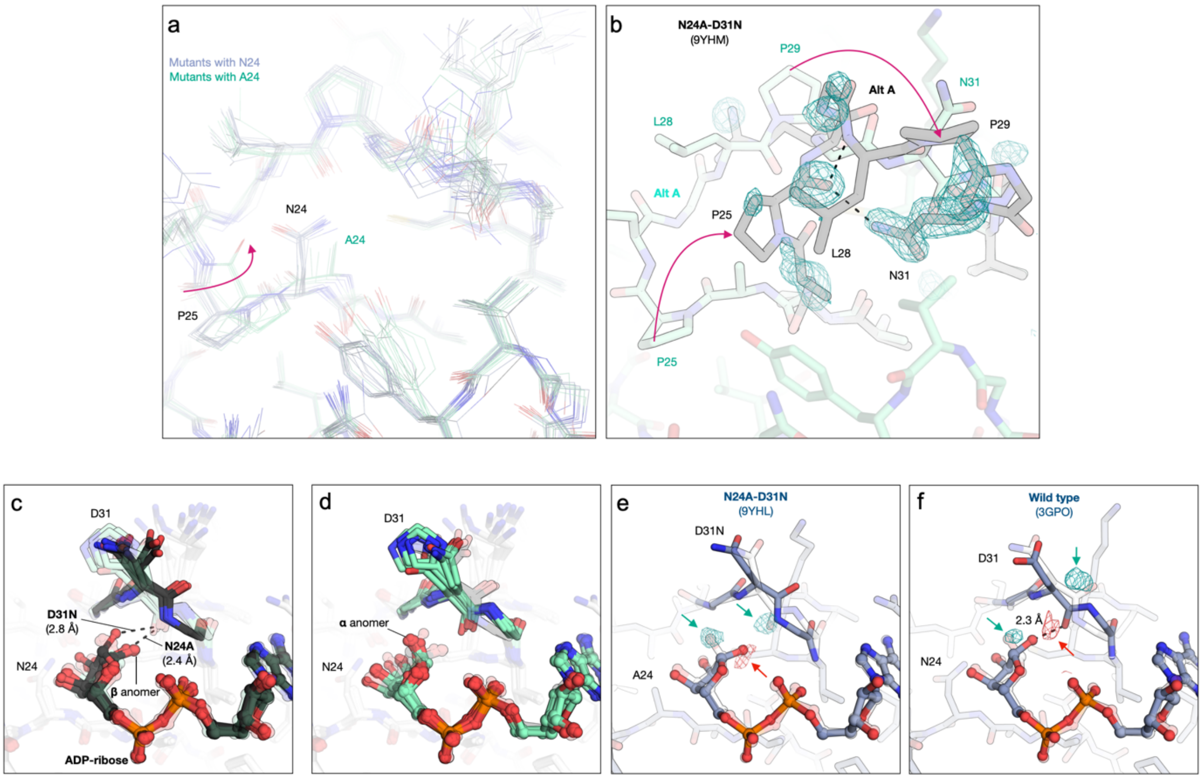


**Supplementary Figure 5. Conformational and composition heterogeneity in the CHIKV nsP3 macrodomain. a**, Alignment of all apo structures colored by the residue at position 24 (asparagine = blue, alanine = green). The peptide flip in P25 is observed in some, but not all, of the structures with A24. **b,** Residues 23-33 undergo a large conformational change in chain D of the N24A-D31N structure (PDB 9YHM). The difference electron density map ($F_O$-$F_C$, 3 σ) prior to modeling the alternative conformation is shown. **c**, Alignment of all CHIKV nsP3 macrodomain mutant structures in complex with ADP-ribose. The β- and α-anomers of ADP-ribose are shown with black and green sticks, respectively. The conformation of residue 31 compatible with the β-anomer is shown with black sticks, while the clashing conformation is shown with green sticks. This pattern was not observed for the structures of N24A (PDB 9YHH, chain D) and D31N (PDB 9YHD, chain D) in complex with ADP-ribose, where the β-anomer binds with the N31 carbonyl in the buried conformation. **d**, Same as **c**, but with inverted transparency. **e**, The difference electron density map ($F_O$-$F_C$, ± 3 σ) is shown prior to modeling the α-anomer of ADP-ribose or the buried conformation of N24A-D31N (PDB 9YHL, chain D). **f**, The difference electron density map ($F_O$-$F_C$, ± 3 σ) calculated from the deposited structure factor amplitudes of a previously reported structure of CHIKV nsP3 macrodomain in complex with ADP-ribose (PDB 3GPO)[34] reveals peaks that are consistent with both ADP-ribose anomers and both D31 conformations.

Supplementary Figure 6

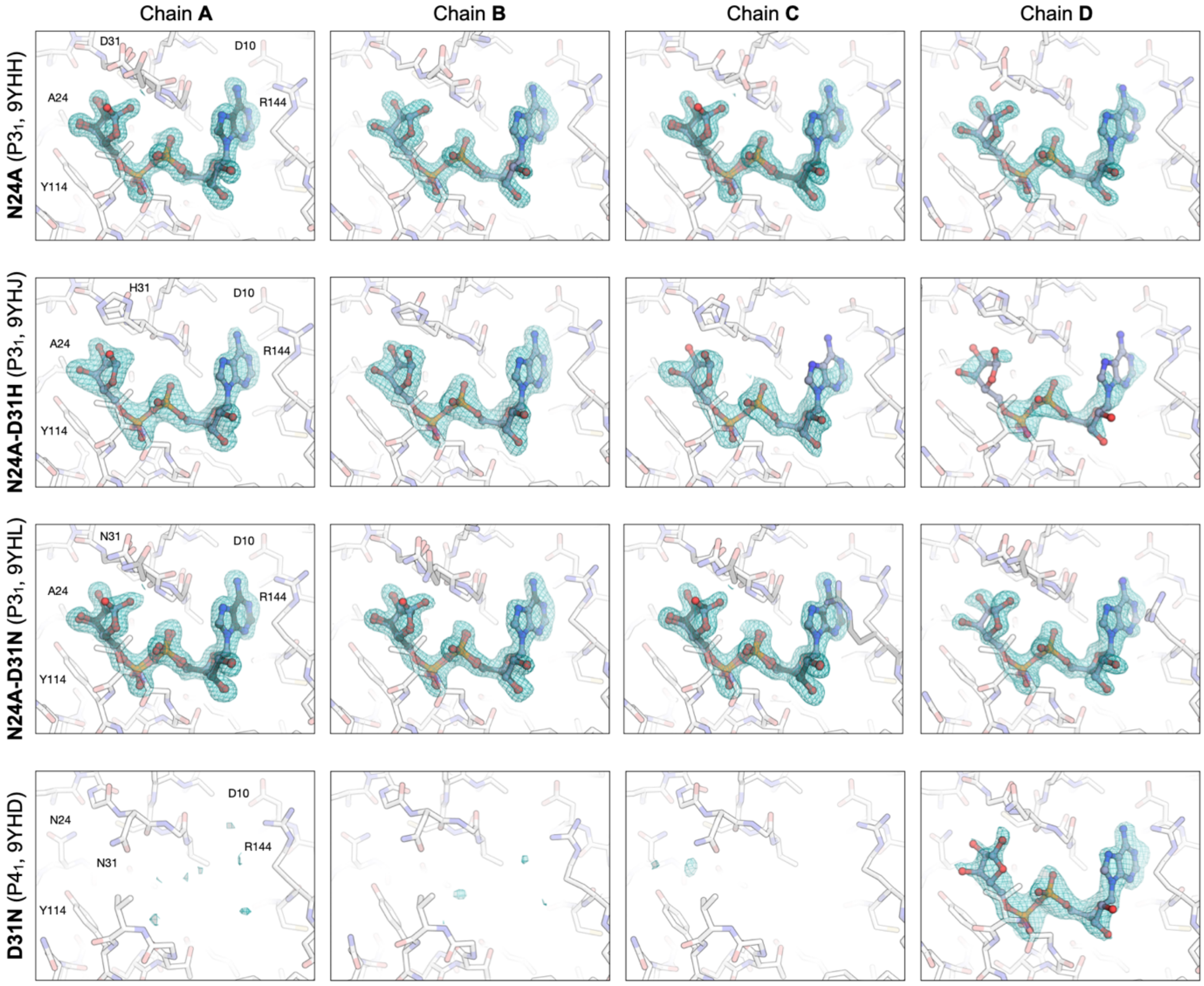


**Supplementary Figure 6. Structures of CHIKV nsP3 macrodomain mutants in complex with ADP-ribose.** Difference electron density maps ($F_O$-$F_C$, 3 σ) calculated prior to ADP-ribose placement for the four structures of CHIKV nsP3 macrodomain determined in complex with ADP-ribose. In the D31N structure, ADP-ribose was only observed in chain D.

**Supplementary Table 1. Macrodomain protein sequences used for multiple sequence alignment and phylogram construction.**

**(Supplementary_Table_1.xlsx)**

**Supplementary Table 2. Sequence analysis of CHIKV macrodomain mutations recovered from viral RNA in individual *Aedes* mosquitoes infected with N24 mutant viruses. (Supplementary_Table_2.xlsx)**

**Supplementary Table 3. Sequence analysis of CHIKV macrodomain mutations recovered from viral RNA in individual *Aedes* mosquitoes infected with D31 mutant viruses. (Supplementary_Table_3.xlsx)**

**Supplementary Table 4. Thermostability and ADP-ribose binding of WT and mutant macrodomains measured by differential scanning fluorimetry (DSF).** Data are reported as mean ± SD for three technical replicates.

| | Tm | + 1mM ADPr | $\Delta T_m$ |
|---|---|---|---|
| WT | 43.2 ± 0.1 | 46.5 ± 0.1 | 3.2 ± 0.1 |
| N24A | 35.9 ± 0.4 | 40.0 ± 0.1 | 4.1 ± 0.4 |
| N24D | 41.3 ± 0.2 | 46.1 ± 0.2 | 4.7 ± 0.2 |
| D31H | 41.9 ± 0.2 | 45.1 ± 0.3 | 3.2 ± 0.4 |
| D31N | 41.3 ± 0.3 | 45.4 ± 0.1 | 4.1 ± 0.3 |
| N24A-D31H | 35.3 ± 0.4 | 39.1 ± 0.6 | 3.8 ± 0.7 |
| N24A-D31N | 36.2 ± 0.4 | 39.4 ± 0.1 | 3.2 ± 0.4 |
| N24D-D31H | 40.7 ± 0.3 | 45.6 ± 0.5 | 4.9 ± 0.6 |
| N24D-D31N | 40.4 ± 0.1 | 45.6 ± 0.2 | 5.1 ± 0.2 |

**Supplementary Table 5. X-ray data collection and refinement statistics. (Supplementary_Table_5.xlsx)**
